# Brain morphological pattern is associated with the presence, severity, and transition of transdiagnostic psychiatric disorders in preadolescents

**DOI:** 10.64898/2026.02.17.706371

**Authors:** Nanyu Kuang, Christopher J Hammond, Betty Jo Salmeron, Xiang Xiao, Danni Wang, Laura Murray, Hong Gu, Tianye Zhai, Hui Zheng, Justine Hill, Maria Scavinicky, Hanbing Lu, Amy Janes, Thomas J Ross, Yihong Yang

## Abstract

Cognitive function, psychological processes, mental states, and behaviors are key dimensions of human subjective experience that separately relate to mental disorders across diagnostic categories. However, whether these dimensions are linked to common or distinct brain morphological patterns that convey risk or resilience for psychiatric disorders remains unclear. The current study is a longitudinal investigation on 11,875 youths from the Adolescent Brain Cognitive Development (ABCD) Study aged 9-10 years at baseline. A machine learning approach based on canonical correlation analysis was used to identify latent dimensional associations of cortical morphology (4 metrics: surface area, cortical and subcortical volume, cortical thickness, and sulcal/gyral depth) with multidomain behavioral assessments including cognitive scores and psychological measures indexing motivation, impulse control, mental states, and behaviors across a normative continuum from healthy to pathological. Across morphological measures, we identified a robust latent brain structural variate that correlated positively with cognitive performance and negatively with psychological measures indexing greater psychology. Notably, higher scores on this brain variate reflected larger cortical surface area and cortical volume—especially in the temporal gyri—together with a posterior-anterior gradient in cortical thickness, showing relatively greater thickness in occipital, parietal, and temporal cortices and lower thickness in cingulate and frontal regions. This brain variate and the related cognitive-psychological-behavioral variate remained stable at the 2-year follow-up, demonstrating temporal consistency. Importantly, the brain variate showed a dose-dependent relationship with the cumulative number of psychiatric diagnoses assessed concurrently and at 2-year follow-up, with lower brain variate scores being associated with higher numbers of comorbid diagnoses. In addition, the brain scores were associated with longitudinal transitions between healthy and diagnosed states over the 2-year study period, in which lower scores at baseline were associated with persistent psychiatric diagnoses whereas higher scores at baseline were associated with persistent healthy states, suggesting that the brain scores capture a vulnerability– resilience continuum for psychopathology. By revealing shared brain structural substrates across conventional diagnostic boundaries, these findings advance the neurodevelopmental understanding of psychiatric disorders and highlight the potential utility of morphology-informed approaches for early screening and intervention in youth.

## Introduction

Psychiatric disorders contribute to significant costs to society in the form of increased suffering, reduced functioning, and increased mortality^1^, yet the lack of established biomarkers to aid diagnosis and treatment has hindered progress in psychiatry^2^. Over the past decades, the focus on diagnostic categories based primarily on clinical symptoms has failed to identify biomarkers specific to these categories potentially due to overlapping symptoms^3^ and high comorbidity among psychiatric disorders^4,5^, suggesting that current diagnostic categories may not necessarily reflect distinct underlying neural mechanisms. Instead, recent large-scale neuroimaging meta-analyses^6-10^, including work from the Enhancing Neuro Imaging Genetics Through Meta Analysis (ENIGMA) Consortium, have provided evidence for shared neurobiological alterations across multiple psychiatric diagnoses. For instance, a transdiagnostic pattern of gray matter loss in the anterior insula and dorsal anterior cingulate was identified^7^, while a shared latent factor for brain structural abnormalities across six major psychiatric disorders was reported^10^. Improved understanding of neurobiological mechanisms that underlie specific clusters of clinical symptoms and span the artificial boundary between psychiatric diagnoses is needed to advance psychiatric nosology.

Within the context of investigating transdiagnostic mechanisms, individual variability in cognitive function, which is often compromised across a wide range of psychiatric conditions^11,12^, offers an additional dimension for understanding mental health and illness. Previous research on cognitive dysfunction in psychiatry has primarily emphasized identifying distinct patterns of impairment associated with specific psychiatric diagnoses (e.g., schizophrenia)^13^. Studies in healthy populations have established a predominantly heritable latent factor, commonly termed the g factor, reflecting general cognitive functioning^14^. A corresponding construct, the p factor, captures the degree of psychopathological manifestations across various diagnostic categories and symptom clusters, theoretically signifying the broad (mal)functioning of emotional and behavioral regulation in clinical and nonclinical samples^15^. Although the relationship between the g factor and the p factor is complex, most studies show an inverse correlation between these two factors^15,16^ as well as their independent associations with functioning in overlapping brain regions, suggesting that they may share underlying neurobiological foundations.

Developmental considerations further refine our understanding of these shared and distinct neural underpinnings. Adolescence constitutes a sensitive period of development marked by dramatic biological, psychosocial, and behavioral changes including gray matter maturation and cognitive development. Notably, many psychiatric disorders emerge before or during adolescence^17-19^. The integration of large-scale population neuroimaging datasets, combined with technological advances in transdiagnostic neuroimaging have begun to uncover neurobiological alterations that may track vulnerability to psychiatric disorders and cognitive activity prior to overt clinical manifestations^20,21^. Among these neurobiological measures, magnetic resonance imaging (MRI)-based cortical morphological metrics—including surface area, cortical thickness, cortical volume, and sulcal depth—exhibit distinct genetic bases and developmental trajectories^22-26^. At the macroscopic level, these cortical features closely mirror simultaneous microstructural changes. For instance, surface area is predominantly influenced by radial glial cell proliferation during early development, which shapes the number of cortical mini-columns and correlates with myelination processes^27^. Cortical thickness, determined by the number of intermediate progenitor cells within each ontogenetic column, is influenced by synaptic pruning and myelination^28^. Sulcal depth is collectively influenced by complex regional differences in cellular proliferation, coupled with axonal tension and other biomechanical mechanisms^29^. The microstructure of the brain undergoes continuous changes throughout the lifespan. From childhood to adolescence, the processes of axonal myelination and synaptic reorganization in brain regions persist, with the synaptic pruning process resulting in the elimination of more than 40% of excitatory and inhibitory synapses^30^. A growing body of evidence also indicates that structural dynamics in cortical organization are closely associated with both psychopathology and cognition in adolescence^31-33^.To elucidate the interrelationships between cognitive and psychological processes as well as their common and divergent neural underpinnings during development, this study aimed to identify latent neurobehavioral relationships between cortical morphological features and a comprehensive set of behavioral measures spanning cognitive, motivational, affective, perceptual, self-regulatory, and behavioral dimensions in preadolescent youth (Fig. 1). We analyzed longitudinal data from the ABCD Study^34,35^, which includes brain structure and comprehensive cognitive and psychological assessments from a large sample of 9-10-year-old children at baseline and at 2-year follow-up. A machine-learning approach, canonical correlation analysis (CCA)^36,37^, was utilized to identify latent components without a priori assumptions^38-40^ from two high-dimensional datasets (cortical morphological features and cognitive-psychological-behavioral measures) that show maximal cross-set correlations. This multivariate approach is particularly powerful for distilling complex, multidomain relationships in large developmental cohorts. Prior applications of CCA in the ABCD Study have already successfully identified shared latent dimensions linking structural brain measures with cognitive and psychological outcomes. For example, a key study evaluating the relationship between cortical morphology and cognition and psychopathology, revealed a major variate indicating larger global brain volume and surface area, alongside localized prefrontal/cingulate thinning, was associated with enhanced cognition and fewer psychological symptoms^41^. Of note, applying a similar approach to resting-state functional connectivity (rsFC) data, our group recently discovered a functional connectivity-based variate that was associated with higher cognitive function and lower psychology processes^42^. Furthermore, recent work validated the stability and reproducibility of CCA-derived brain–behavior relationships within the ABCD cohort, reinforcing the robustness of this analytical method for identifying reliable neurodevelopmental markers^43^. The current study extends these lines of research to evaluate a more comprehensive set of cortical morphological features and provides a longitudinal assessment of their clinical relevance. Specifically, the present study uses CCA to (1) derive latent dimensions linking cortical morphology with multidomain cognitive and psychological measures, (2) examine their longitudinal stability over a 2-year follow-up, and (3) evaluate their clinical relevance by testing associations with the burden and course of psychiatric diagnoses in youth.

**Fig. 1.**
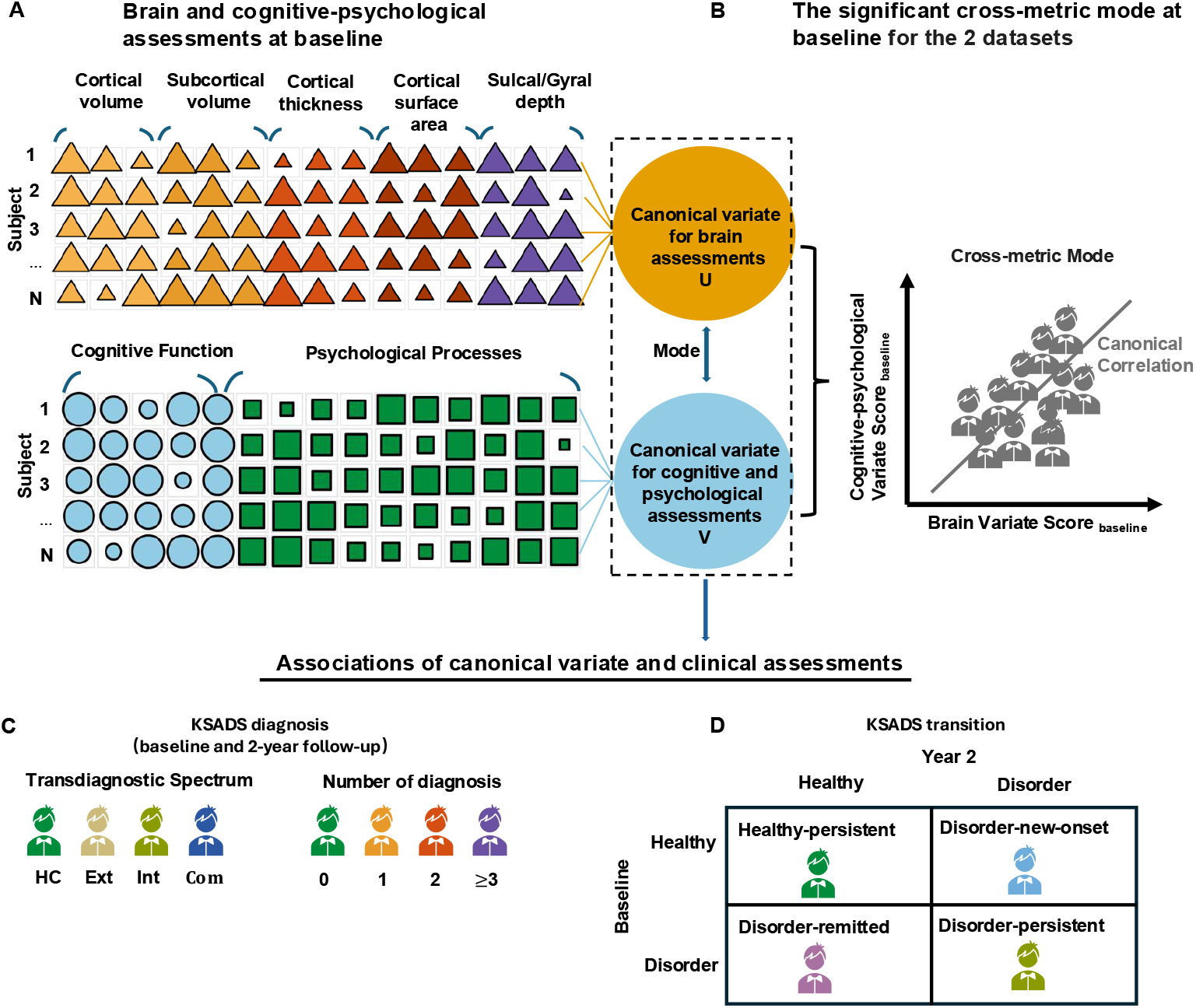
Schematic workflow of the study. (A) Brain and cognitive-psychological assessments at baseline; (B) Significant cross-metric mode at baseline identified by canonical correlation analysis (CCA); (C) The associations of canonical variate and KSADS diagnoses at baseline and 2-year follow-up; (D) The associations of canonical variate and diagnostic status transitions over two years. Abbreviations: HC: healthy control; Ext: externalizing disorders; Int: internalizing disorders; Com: comorbidity between externalizing and internalizing disorders; KSADS: Kiddie Schedule for Affective Disorders and Schizophrenia for DSM-5.

## Results

### Participants, Brain Morphological Measures, and Cognitive and Psychological Assessments

Of 11,985 participants from the ABCD study, 8,672 at baseline and 6,218 at 2-year follow-up were included in the CCA analyses based on quality control of their MRI data and availability of their cognitive-psychological assessments (See Supplementary Information and Fig.S1 for more details). Demographic information of the included participants is summarized in Table S1.

The cortical brain was segmented into 68 regions using the Desikan-Killiany atlas^44^. Brain morphological metrics (surface area, cortical volume, cortical thickness, and sulcal/gyral depth) were extracted from these cortical regions. Additionally, these morphological metrics were obtained from the Destrieux atlas^45^ (148 cortical regions) to further support our findings. Subcortical volumes of the bilateral thalamus, caudate, putamen, pallidum, hippocampus, amygdala, accumbens, and ventral diencephalon (a total of 16 volumes) were computed based on the Fischl atlas.

Cognitive assessments included 13 cognitive scores indexing general cognition and domain-specific cognitive functioning derived from participant performance on 10 neuro-cognitive tests^46^ (e.g., NIH Toolbox Cognition Battery) and 3 neuroimaging tasks^47^. Psychological assessments included 31 psychological variables indexing approach-vs-avoidant motivational tendencies, impulse control, and emotional, thought, perceptual, somatic, attentional, and conduct problems derived from youth and parent/caregiver responses on the Behavioral Inhibition System/Behavioral Activation System (BIS/BAS) Scale (youth-report)^48,49^, Urgency, Premeditation, Perseverance, Sensation Seeking (UPPS-P) Impulsive Behavior Scale (youth-report)^50,51^, Child Behavioral Checklist (CBCL, parent-report^52^), Parent General Behavior Inventory-10 Item Mania Scale^53^ (PGBI-10, parent-report^54^), and Psychosis Prodrome Questionnaire-Brief (PQ-B, youth-report^55^). Measurements, variables, and abbreviations of cognitive and psychological assessments are listed in Table S2 and S3, respectively.

### Latent Dimensions Linking the Brain Structure with Cognitive Function and Psychological Processes

We performed a cross-metric analysis—which first concatenated surface area, cortical and subcortical volume, cortical thickness, and sulcal/gyral depth into a single brain-morphology matrix and then applied CCA to relate this unified dataset to the cognitive-psychological measures (Fig.S2). This analysis identified one significant brain structural and cognitive-psychological association within the training sets, which generalized to the holdout test sets when compared with a permutation-generated null distribution (Fig. S3). The significant cross-metric CCA mode on Desikan–Killiany atlas showed a correlation of ρ = 0.32 (familywise error rate–adjusted permutation test p_fwer_ < 10^−5^) in the training set and ρ = 0.30 (p_fwer_ < 10^−5^) in the test set, as illustrated by the permutation-based correlation distributions in Fig. S3, and uniquely accounted for a large proportion of redundant variance^56^ between the brain structural and cognitive-psychological datasets. We also performed the same analyses on the Destrieux atlas^45^ (Fig S4); the significant brain structural and cognitive-psychological associations identified in this parcellation closely resemble those observed with the Desikan–Killiany atlas in both brain structural loadings and cognitive-psychological loadings. Accordingly, we present only results with the Desikan–Killiany atlas in the main text.

To evaluate how different morphological metrics and cortical territories shaped the canonical brain variate, we ran a two-way ANOVA with Metric (surface area, cortical volume, cortical thickness, sulcal/gyral depth) and Region (occipital, parietal, temporal, insula, cingulate, frontal; Fig. 2A-C) as factors. This analysis revealed significant main effects of Metric (F = 54.36, p < 10^−5^) and Region (F = 16.09, p < 10^−5^), as well as significant Metric × Region interaction effects (F = 5.01, p < 10^−5^), indicating that the cross-metric brain variate is not a uniform “size” factor but is differentially expressed across indices and anatomical territories. Post-hoc Tukey tests confirmed that surface area and volume showed larger mean loadings than cortical thickness and sulcal/gyral depth (FDR-corrected p < 10^−3^). Mapping the loadings onto the cortical surface (Fig. 2D-G) highlighted two key spatial signatures. First, surface area and cortical volume exhibited broadly similar patterns, with the highest positive loadings concentrated in temporal cortices (e.g., middle and inferior temporal gyri). Second, cortical thickness showed a clear posterior–anterior gradient, with positive loadings most evident in occipital, parietal and temporal cortices and negative loadings in cingulate and lateral frontal regions. Sulcal/gyral depth made comparatively weaker contributions to the canonical pattern, with mainly negative loadings in cingulate cortices. Subcortical volumes loaded positively overall, with the strongest contributions in the left hippocampus and left ventral diencephalon.

**Fig. 2.**
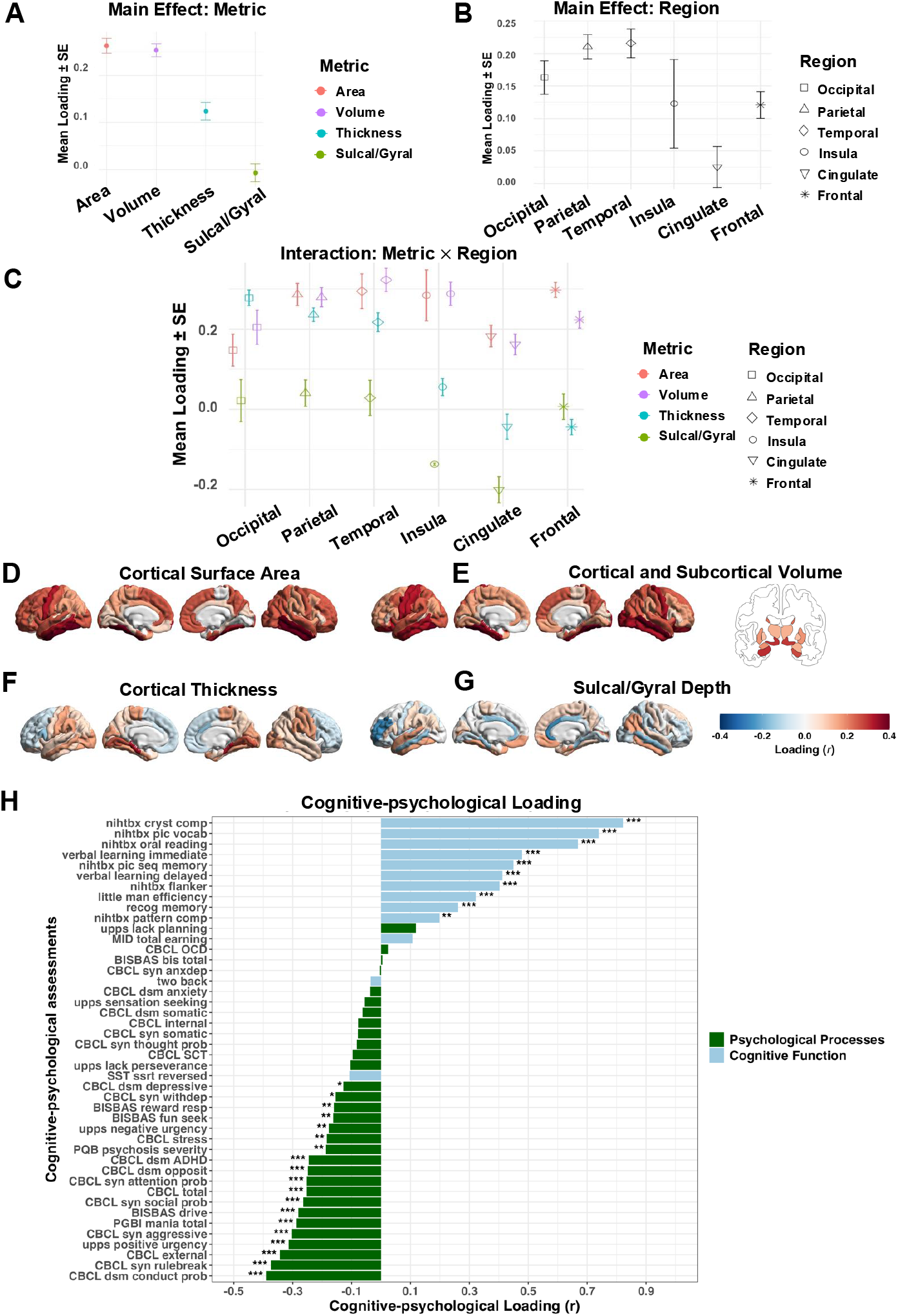
The cross-metric mode based on canonical correlation analysis (CCA). (A) Two-way ANOVA – main effect of morphological Index, (**B**) anatomical Region and (C) Index×Region interaction on loadings. The whole cross-metric brain variate loadings, mapped onto four individual metrics: (D) surface area, (E) cortical and subcortical volume, (F) sulcal/gyral depth and (G) cortical thickness. (H) Loading of each cognitive and psychological assessment on the cognitive-psychological variate. Abbreviations are listed in Tables S2 and S3. False discovery rate– adjusted p values: ∗p <.05, ∗∗p <.01, ∗∗∗p <.001.

Fig.2H shows the significant loading of each cognitive and psychological measure on the latent cognitive-psychological variate of the cross-metric mode. Nearly all assessments related to cognitive function showed positive loadings on the latent behavior variate. In contrast, most psychological assessments, including those indexing approach motivation (BIS/BAS drive), poorer impulse control (UPPS-P U and PU scales), and emotional, perceptual, thought, and attentional problems (CBCL subscales), showed negative loadings. Among these cognitive measures, the Crystallized Composite Score demonstrates the highest loading on the latent behavior variate (r = 0.82). Furthermore, differences were observed across externalizing and internalizing domains of psychological processes. Measures indexing the externalizing spectrum problems generally exhibited higher loadings (e.g., r = −0.37 for CBCL Rule Breaking and r = −0.39 for CBCL Conduct Problem) compared to measures indexing the internalizing spectrum problems (e.g., r = −0.0035 for CBCL Anxious-Depression and r = −0.03 for CBCL dsm_anxiety); Statistical significance was assessed using bootstrapping (5000×), false discovery rate–corrected *p* <.05.

### Rank-order stability of the Brain Structural Variates in Longitudinal Assessments

To determine whether the brain structural variate identified in our main cross-sectional CCA analysis shows temporal stability (i.e., is stable over time), we performed longitudinal validation tests using follow-up data collected 2-years later. Specifically, we projected the cross-metric mode estimated from the baseline training set (brain structural data) onto a test dataset consisting of brain structural assessments at the 2-year follow-up (see Supplementary Information and Fig. S6). The resulting baseline brain structural variate scores were highly correlated with their 2-year counterparts (r = 0.91), indicating strong rank-order stability—that is, individuals with high (or low) brain variate scores at baseline tended to continue to have high (or low) scores two years later, and their relative positions along the brain–cognitive–psychological axis were largely preserved (Fig. S7). To further examine age- and diagnosis-related effects, we fit a linear mixed-effects model based on the brain variate scores from both baseline and the 2-year follow-up for all participants, modelling the brain variate as a function of age and diagnosis (healthy plus the ten diagnostic groups shown in Fig. 3). This analysis revealed a small but significant age-related increase in brain variate scores across the cohort (F = 25.88, p = 3.7 × 10^−7^), whereas the age-by-diagnosis interaction was not significant (F = 1.59, p = 0.10). Taken together, these results indicate that the brain structural variate shows high rank-order stability at the individual level and broadly parallel developmental trajectories across diagnostic groups, with only modest age-related shifts in mean level over the 2-year interval.

**Fig. 3.**
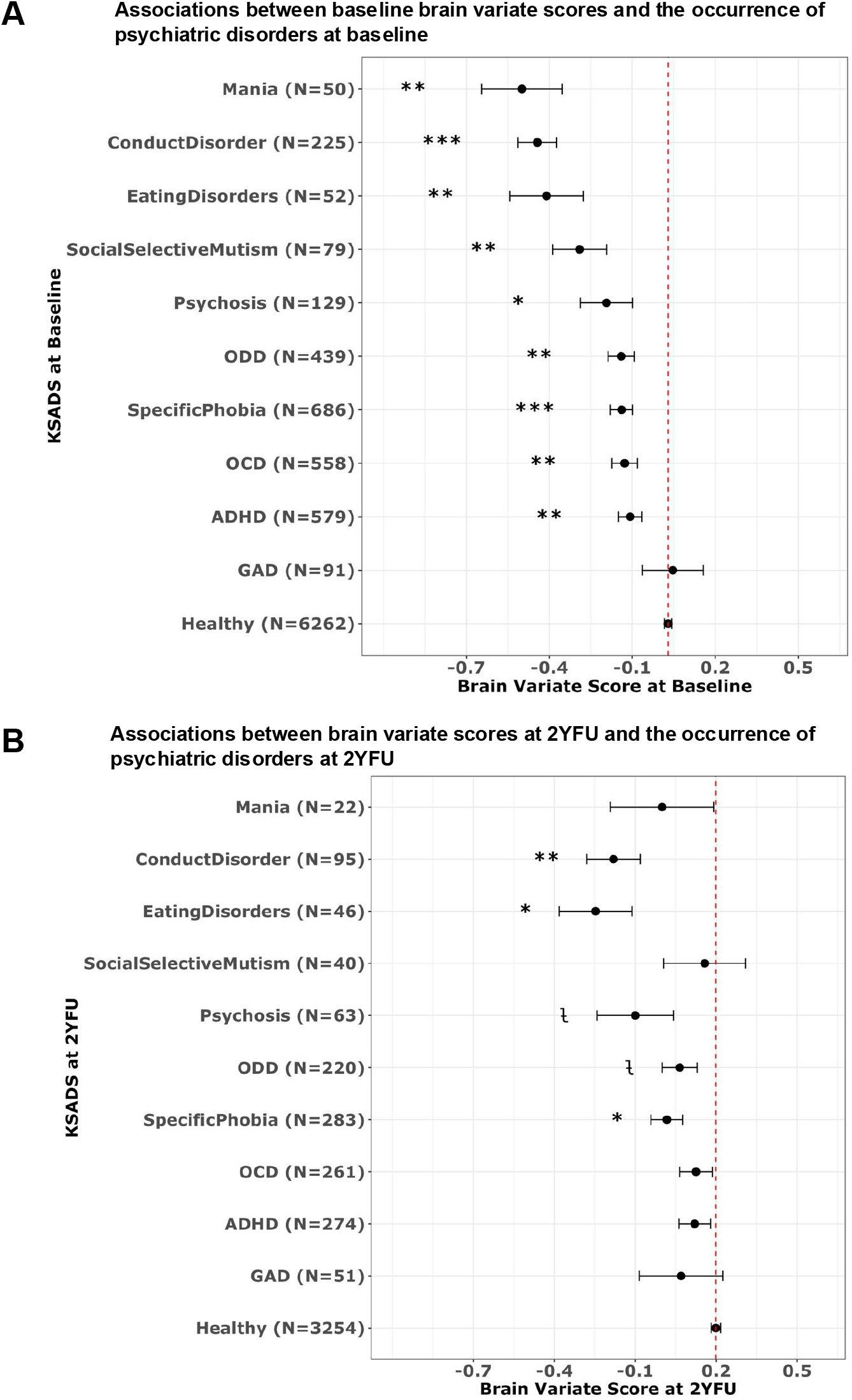
Relationships between brain variate score and the occurrence of psychiatric disorders at baseline and the 2-year follow-up. (A) Case-control analyses comparing brain variate scores at baseline between specific diagnosis and healthy controls at baseline. (B) Case-control analyses comparing brain variate scores at 2-year follow-up between specific diagnosis and healthy controls at 2-year follow-up. Error bars indicate standard error. T tests were used to compare differences between each diagnostic subgroup to the healthy controls. Ł: p unadjusted <.05; False discovery rate–adjusted p values: *: p adjusted <.05; **: p adjusted <.01; ***: p adjusted <.001. The full names of each diagnostic abbreviation are provided in the Supplementary Information abbreviation list.

### Consistency of the Cross-metric Mode Across Cognitive and Psychological Domains

To characterize the stability from cognitive and psychological processes to the cross-metric brain structural variate cross-metric mode, we conducted supplemental analyses re-running the CCA procedure twice, first while holding out the cognitive measures (to determine the variance in cross-metric mode that was uniquely related to psychological processes) and second holding out the psychological measures (to determine the variance in cross-metric mode that was uniquely related to cognition). The brain structural variates showed very high consistency among the 3 cases (combined vs. cognition only, r = 0.95; combined vs. psychological processes only, r = 0.81), see Fig.S8-S9.

### Association between Brain Structural Variate Score and Specific Psychiatric Diagnoses

To examine disorder-specific relationships, all participants were grouped by their Kiddie Schedule for Affective Disorders and Schizophrenia for DSM-5 (KSADS-5) diagnoses at baseline and 2-year follow-up, and the brain structural variate at baseline and 2-year follow-up of each diagnostic group was compared with the group with no current psychiatric diagnosis using Welch two-sample t-tests.

Individuals with various current psychiatric disorders at baseline and at the 2-year follow-up exhibited lower brain structural variate scores at each time point (see Fig. 3). Specifically, compared with healthy control participants, individuals diagnosed at baseline with mania, conduct disorder, eating disorders, social selective mutism, psychosis, oppositional defiant disorder (ODD), specific phobia, obsessive–compulsive disorder (OCD) and attention deficit hyperactivity disorder (ADHD) exhibited significantly lower brain variate scores at baseline (see Fig. 3A). At the 2-year follow-up, this pattern persisted for those with conduct disorder, eating disorders and specific phobia (see Fig. 3B). All findings remained significant after correction for multiple comparisons using the false discovery rate (see Tables S4-S5).

### Association between Brain Structural Variate Score and Cumulative Number of Psychiatric Diagnoses

We examined associations between the brain structural variate score and cumulative number of diagnoses at baseline and 2-year follow-up. Participants who had both brain, cognitive-psychological and clinical measures available at baseline and 2-year follow-up were stratified into four groups (0, 1, 2 and ≥3) based on the cumulative number of current psychiatric disorders assessed via KSADS interview at baseline (0 (N = 6262), 1 (N = 1211), 2 (N = 414) and ≥3 (N = 252)) and 2-year follow-up ((0 (N = 3254), 1 (N = 655), 2 (N = 186) and ≥3 (N = 104))). Consistently, lower brain structural variate scores were associated with greater cumulative number of psychiatric disorders, regardless of type (Figs. 4A-B and Tables S6-S7). Further, many of these associations exhibited dose-response relationships whereby children with more psychiatric diagnoses had lower brain structural variate scores, and vice versa.

**Fig. 4.**
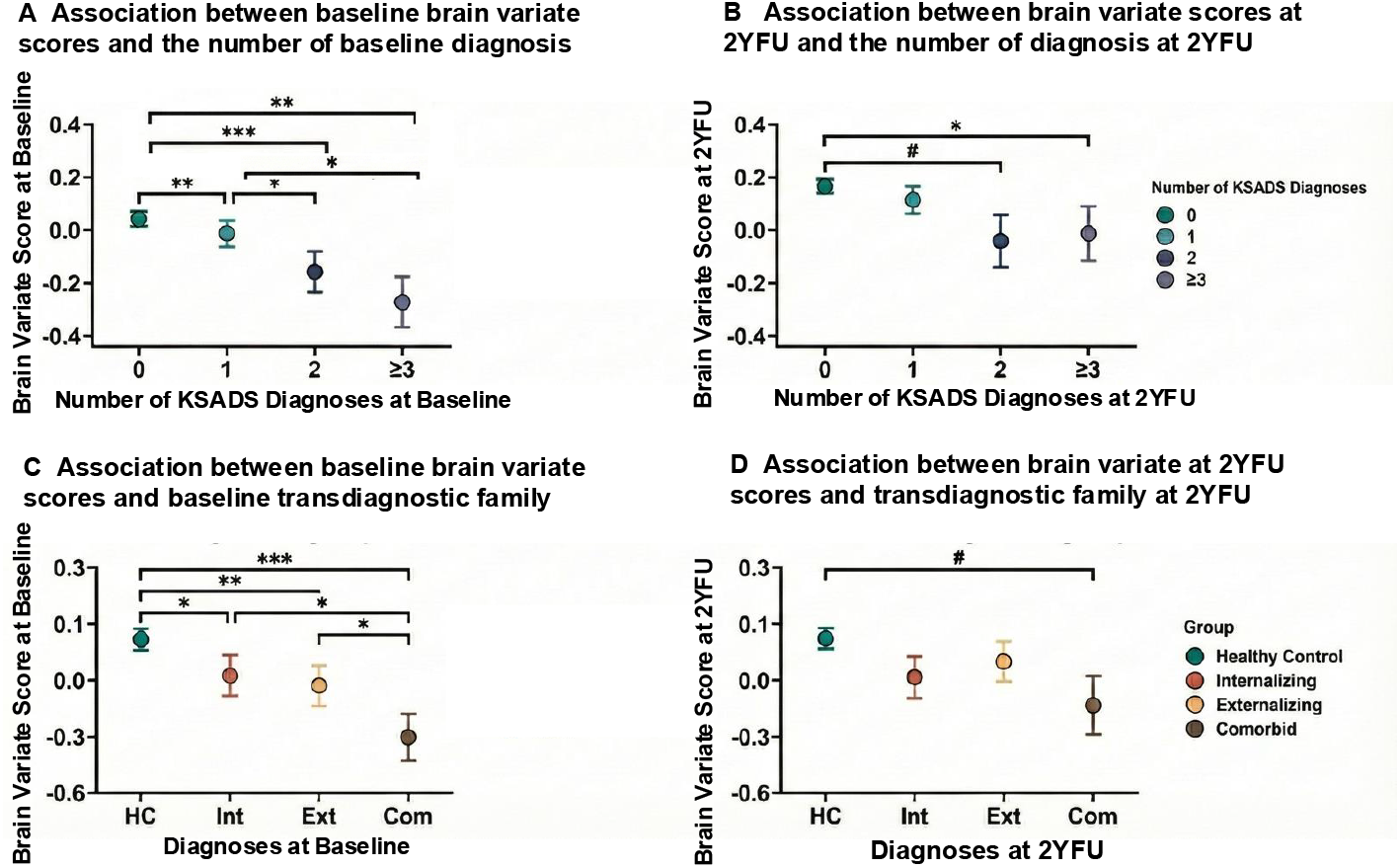
Relationships between brain variate score and transdiagnostic disorders. **(A)** Group-wise comparison of brain variate score at baseline by cumulative number of comorbid disorders at baseline. (B) Group-wise comparison of brain variate score at 2-year follow-up (2YFU) by cumulative number of comorbid disorders at 2-year follow-up. (C) Group-wise comparison of brain variate score at baseline among the following groups at baseline: internalizing disorders (Int), externalizing disorders (Ext), comorbid (Com; internalizing + externalizing) disorders, and healthy controls (HC). (D) Group-wise comparison of brain variate score at 2-year follow-up among the following groups at 2-year follow-up: Int, Ext, Com, and HC. Dots and lines in the plot indicate mean and standard error of the variate score in each group. Pairwise group differences were post hoc compared using the Games-Howell procedure. #: p _unadjusted_ <.05; *: p _adjusted_ <.05; **: p _adjusted_ <.01; ***: p _adjusted_ <.001.

Specially, One-way ANOVAs testing differences in brain structural variate scores across groups with different numbers of psychiatric disorders found evidence of concurrent and prospective associations between brain structural variate scores and cumulative number of psychiatric disorders. ANOVAs on brain structural variate scores at baseline and KSADS diagnoses at baseline revealed a significant effect of brain structural variate (F = 16.71, p <.00001; see Fig. 4A). ANOVAs on brain structural variate scores at 2-year follow-up and KSADS diagnoses at 2-year follow-up were also significant suggesting prospective associations between brain structural variate scores and cumulative number of psychiatric disorders (F = 4.34, p <.01, see Fig. 4B).

### Association between Brain Structural Variate Score and Transdiagnostic Spectrum

To investigate the relationship between the canonical brain structural variate score and broad transdiagnostic diagnostic families at both baseline and two-year follow-up, based on the grouping framework of transdiagnostic diagnostic families^25,57^, participants who had both brain, cognitive-psychological and clinical measures available at baseline and 2-year follow-up were stratified into two overarching diagnostic spectra: externalizing disorders (N_baseline_ = 707,N_2Y_ = 368)—comprised of attention-deficit/hyperactivity disorder, oppositional defiant disorder, and conduct disorder—and internalizing disorders (N_baseline_ = 505,N_2Y_ = 271) —including dysthymia, major depressive disorder, disruptive mood dysregulation disorder, agoraphobia, panic disorder, specific phobia, separation anxiety disorder, social anxiety disorder, generalized anxiety disorder, and post-traumatic stress disorder. Based on these definitions, study participants were stratified into four mutually exclusive cohorts for comparative analysis: healthy controls, externalizing disorders, internalizing disorders, and comorbid externalizing + internalizing disorders (N_baseline_ = 234, N_2Y_ = 90). Across both time points, brain structural variate scores were lower as the number of transdiagnostic spectra increased—healthy controls showed the highest scores, followed by the externalizing and internalizing groups (which did not differ significantly from each other), with comorbid cases exhibiting the lowest scores—indicating that greater transdiagnostic burden, rather than diagnostic category per se, associates reductions in the canonical brain variate (Figs. 4C-D).

ANOVAs using brain structural variate scores at baseline and transdiagnostic spectrum at baseline revealed a significant group effect (F = 10.80, p <.00001; see Fig. 4C). The result showed that each disorder group—externalizing and internalizing disorders and their comorbidity exhibited significantly lower variate scores than healthy controls, and the comorbid group scored even lower than both the externalizing and internalizing disorders. However, there was no significant difference between the externalizing and internalizing disorders. ANOVAs using brain structural variate scores at 2-year follow-up and transdiagnostic spectrum at 2-year follow-up also demonstrated a significant group effect and similar pattern (F = 2.79, p <.05; see Fig. 4D).

### Association between Brain Structural Variate Score and Psychiatric Diagnostic Status Transitions from Baseline to 2-Year Follow-up

We sought to determine whether brain structural variate scores track changes in diagnostic status over a 2-year interval. To this end, based on whether the participants had psychiatric diagnoses at baseline and 2-year follow-up, participants were categorized into 4 subgroups: Healthy Persistent (0 psychiatric diagnosis at both baseline and 2-year follow-up, N = 5082), Disorder Remitted (≥1 psychiatric diagnosis at baseline and 0 psychiatric diagnosis at 2-year follow-up, N = 893), Disorder New Onset (0 psychiatric diagnosis at baseline and ≥1 psychiatric diagnosis at 2-year follow-up, N = 702) and Disorder Persistent (≥1 psychiatric diagnosis at both baseline and 2-year follow-up, N = 893). Using one-way ANOVAs, we compared each group’s brain structural variate scores at baseline and their longitudinal change (brain structural variate scores at 2-year follow-up minus brain structural variate scores at baseline).

The brain structural variate scores—as well as their longitudinal change—showed a significant association with transition group membership (See Fig. 5 and Tables S8-S9). At baseline, the brain structural variate score differed significantly across transition groups (F = 7.26, p <.0001, see Fig. 5A). Furthermore, the longitudinal change in brain structural score differed significantly across groups (F = 5.01, p <.01, see Fig. 5B). Specifically, at baseline, the Healthy Persistent group showed the highest brain variate score, followed by the Disorder New Onset, Disorder Remitted, and Disorder Persistent groups. From baseline to 2-year follow-up, the Healthy Persistent group exhibited largest positive changes in brain variate score, followed by the Disorder New Onset and Disorder Remitted groups, while the Disorder Persistent group demonstrated negative changes (on average) in the brain score. These findings highlight the differential longitudinal trajectories in brain structure among clinical subgroups, underscoring the potential of brain structural variates as sensitive markers of clinical status transitions.

**Fig. 5.**
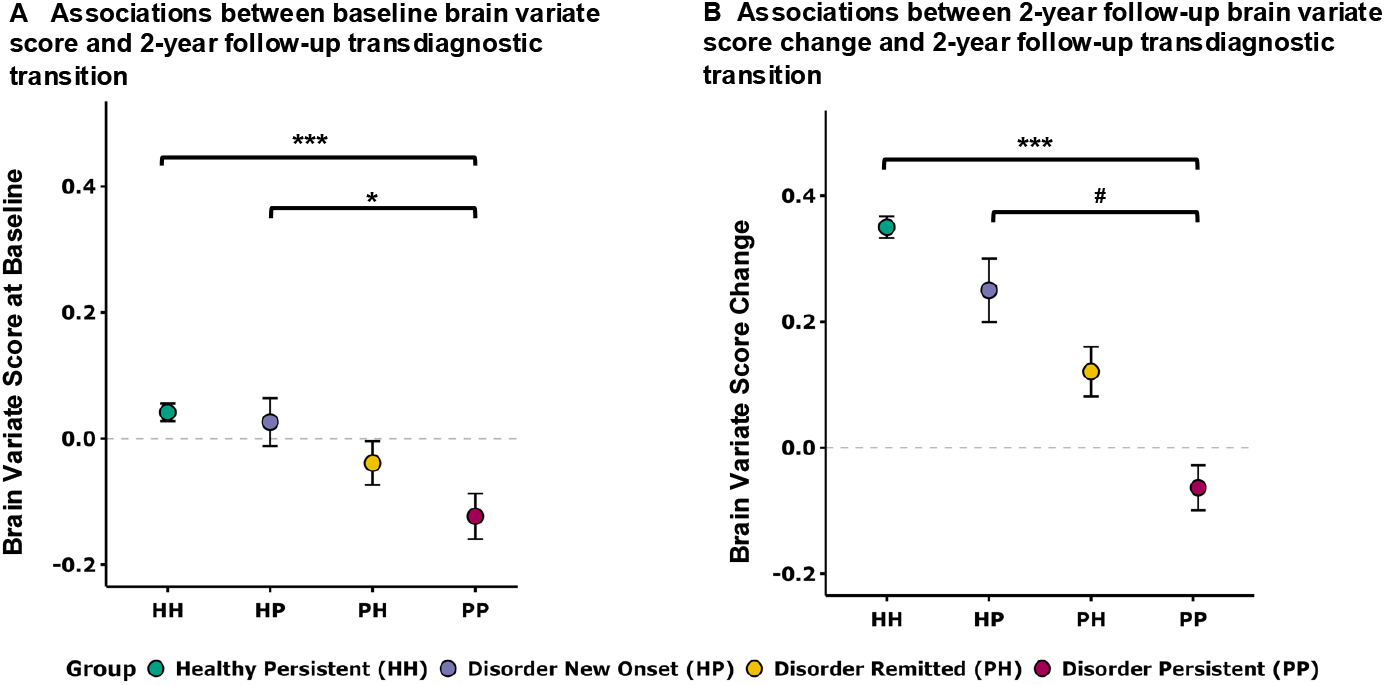
Relationships between brain variate score and the longitudinal transition of Kiddie Schedule for Affective Disorders and Schizophrenia for DSM-5 (KSADS-5) diagnosis status. (A) Groupwise comparison of brain variate score at baseline (A) and change in brain variate score (B) in 4 groups defined by their diagnostic transition. Dots and lines in the plot indicate mean and standard error of the variate score in each group. Pairwise group differences were post hoc compared using the Games-Howell procedure. #: p _unadjusted_ <.05; *: p _adjusted_ <.05; **: p _adjusted_ <.01; ***: p _adjusted_ <.001. HealthyPers = Healthy Persistent group, DisorderNewOnset = Disorder New Onset group, DisorderRemit = Disorder Remitted group, DisorderPers = Disorder Persistent group.

## Discussion

Applying CCA to a multidimensional dataset that included four structural MRI-derived brain morphology metrics (surface area, cortical thickness, cortical and subcortical volume, and sulcal/gyral depth) and comprehensive measurements of cognition and psychological processes from 8,672 US preadolescents with baseline data from the ABCD study, we identified a single latent brain structural variate that covaried with a broad array of cognitive and psychological measures. The main cross-metric variate that combined all four morphological features showed the greatest reliability and explained the largest amount of variance in behavioral domains; however, some cross metric differences in regional loading patterns were observed. Cortical thickness exhibited a posterior–anterior gradient, with positive loadings in occipital, parietal, and temporal cortices and negative loadings in cingulate and frontal regions, whereas surface area and cortical volume showed consistent positive loadings—particularly in the temporal gyri. The brain structural variate was positively associated with performance on multiple cognitive tasks and negatively associated with self- and parent-report measurements indexing psychological problems across different domains and categories, respectively. Notably, the brain variate could be robustly identified from either cognitive or psychological domains and showed good rank stability over time. The brain variate showed a dose-dependent relationship with the cumulative number of psychiatric diagnoses assessed concurrently and at 2-year follow-up, with lower brain variate scores being associated with higher numbers of comorbid diagnoses. Finally, the brain variate was associated with longitudinal transitions between healthy and psychiatrically diagnosed states over the 2-year study period. Our results suggest that this latent brain structural variate may represent a unified dimension that captures individual-level variation in transdiagnostic risk for psychology across a vulnerability– resilience continuum during early adolescence.

### Posterior-Anterior Gradient of Cortical Thickness in the Brain Variate

A defining feature of this brain structural variate is a distinct posterior–anterior gradient observed in cortical thickness. Specifically, the variate is characterized by positive loadings (thicker cortex) in posterior regions (temporal, parietal, occipital regions) and negative loadings (thinner cortex) in anterior regions (cingulate and lateral frontal regions). When related to both cognitive and symptom (e.g., CBCL) measures, we found that less thickness in frontal lobe and greater thickness in temporal, parietal, and occipital regions was associated with better cognitive performance and lower scores on psychological measures, particularly externalizing problems. Mechanistically, cortical thinning during late childhood and adolescence reflects ongoing synaptic pruning^58,59^ and myelination^60,61^, processes integral to refining network efficiency^62-64^. Moreover, myelination alters T1-weighted gray-white matter contrast^65^, shifting the gray/white border classification and thereby reducing measured cortical thickness^66^. Primary sensory cortices (ages 4-8) and parietal areas (ages 11-13) complete these maturational processes earlier^67^, and thus greater residual thickness likely indexes preserved synaptic density and dendritic complexity that support superior perceptual integration and cognition but may also predispose to heightened impulsivity and externalizing behaviors^68^. In contrast, frontal lobe completes these maturational processes in later stages of pubertal development^67^, as more complex abilities such as language, attention, and executive functioning emerge^69^. Lower thickness in frontal lobe marks advanced network specialization underpinning executive control yet may weaken inhibitory resilience, manifesting as increased impulsivity and aggression. This pattern is also consistent with previous studies, in which cortical thinning in frontal lobe during adolescence has also been associated with better performance on individual cognitive tasks, including visual^70^, verbal memory^71^, verbal fluency^72^, and executive functioning^73^. Taken together, these findings are suggestive of a developmental heterochrony in which region-specific cortical maturation trajectories map onto both adaptive (enhanced cognition) and maladaptive (externalizing) outcomes.

### Global Contributions of Surface Area and Volume to the Brain Variate

Complementing the gradient observed in thickness, cortical volume and surface area contributed to the brain structural variate through a broadly positive loading pattern across the brain, particularly in the temporal lobe. Larger cortical volume and surface area were consistently associated with higher cognitive performance and lower externalizing problems. This aligns with prior literature linking global surface area to general intellectual ability^74,75^ and negatively to the *p*-factor (general psychology)^76-79^. Our results showing that surface area and cortical thickness exhibit distinct loading patterns (global positive vs. posterior-anterior gradient) within the same latent variate is consistent with the radial unit hypothesis, which posits that these traits are driven by dissociable genetic and cellular mechanisms (progenitor cell proliferation vs. neurogenic division) ^27 79^. The cross-metric variate thus succeeds in integrating these distinct biological pathways into a single, comprehensive phenotype of neural integrity.

### The Variate as a ‘Candidate’ Transdiagnostic Psychology Biomarker

Crucially, this brain structural variate demonstrates utility as a candidate transdiagnostic biomarker. We observed a robust dose-response relationship: lower brain variate scores were associated with a higher cumulative number of psychiatric diagnoses, regardless of the specific disorder type. Our results are consistent with other studies showing common neural alterations across diagnostic categoriesa and align with the Research Domain Criteria (RDoC) framework^80,81^, moving beyond disorder-specific neural alterations to identify core associations spanning diagnostic categories. The brain variate’s non-specific association with a wide array of psychological measurements covering different categories of psychological problems and psychiatric diagnoses suggests it indexes a general susceptibility to psychology. Furthermore, this marker showed high rank-order stability (r = 0.91) over the two-year follow-up. Importantly, it was sensitive to longitudinal clinical changes: individuals who were healthy (i.e., diagnosis free) at baseline and remained healthy at the 2-year follow-up showed the highest scores and positive developmental change, while those who had psychiatric disorders at baseline and continued to have psychiatric disorders at the 2-year follow-up (i.e., persistent disorders) showed the lowest scores and negative change. This indicates the variate is not merely a static trait but tracks the dynamic trajectory of mental health across a continuum of resilience versus risk.

### Integration of Cognition and Psychology

Our findings also provide a neural basis for the intersection of the general cognition factor and the general psychology factor^82-84^. Traditionally viewed as separate constructs, we found that cognitive abilities and psychology risk loaded on opposite ends of the same dimensional behavior variate. The brain structural variate mapped onto this axis, suggesting that the neural architectures supporting high cognitive function are intrinsically linked to those conferring emotional and behavioral resilience. This integration offers a more holistic view of brain-behavior relationships in preadolescence.

## Conclusion

We identified a single latent brain structural variate that links multimodal cortical morphology with cognitive performance and psychology in preadolescence. This variate was robust across cognitive and symptom domains, showed high rank-order stability over two years, and tracked both cumulative diagnostic burden and longitudinal diagnostic transitions. These results support the interpretation of this dimension as a candidate transdiagnostic neural marker of vulnerability-resilience, meriting further prospective and replicative investigation.

## Limitations

This study had some notable limitations. As a secondary analysis of ABCD Study data, the current study relied on the methodological choices related to design and measurement that were made by the primary study team. For example, measures of psychology were mostly parent-reported at this time point, and the findings need to be cross-validated with multi-informant assessments from subsequent data waves. While our report emphasizes transdiagnostic psychology, it is important to note that most study participants scored within the normal range on both psychopathological and cognitive assessments and did not meet criteria for psychiatric diagnoses at either time (assessment) point. However, by targeting resilience enhancement across the full continuum of risk—from health through high-risk states—this analytic framework enables a more nuanced understanding of the processes that underlie vulnerability. Future application of this approach to clinical cohorts characterized by more severe and heterogeneous psychology will be critical for assessing the generalizability, pathophysiological specificity, and translational relevance of these brain-behavior associations, ultimately guiding the refinement of targeted intervention strategies.

## Methods

### Participants

Data from the ABCD project of the National Institute of Mental Health National Data Archive (NDA) release 5.1 (https://wiki.abcdstudy.org/release-notes/start-page.html), includes a cohort of 11,875 youths recruited at baseline from 21 sites across the United States.

### Neuroimaging and Preprocessing

We used post-processed structural MRI and data from the ABCD 5.1 release. Adolescents who met the quality control criteria were included in the imaging-related analyses (imgincl_t1w_include = 1). The detailed preprocessing and quality control procedures have been described previously^85^. The cortical brain was segmented into 68 regions (brain volume, cortical thickness, sulci/gyri depth and surface area) using the Desikan-Killiany atlas^44^ and 16 subcortical volumes were used from the Fischl atlas^86^. Subcortical volumes included the bilateral thalamus, caudate, putamen, pallidum, hippocampus, amygdala, accumbens, and ventral diencephalon. Additionally, the Destrieux atlas^45^ was also used to further support our findings (148 brain regions). For sulci/gyri depth, regions that moved outward during inflation were positive and represent the depths of sulci, and regions that moved inward were negative and represent the height of gyri^87^.

### Assessments of Cognitive Functioning and Psychological Processes

The cognitive-psychological dataset comprised multisource data assessing cognitive functioning and psychological processes across multiple domains. Participants’ cognitive functioning was assessed using performance data from a 10-domain neurocognitive test battery^88^ and behavioral results from 3 neuroimaging tasks^89^ (see Supplementary Information for details). These assessments yielded 13 continuous measures of cognition were obtained per participant. Psychological processes at baseline and follow-up were assessed using multi-informant questionnaire response data from the BIS/BAS, UPPS-P, CBCL, PGBI-10, and PQ-B using validated total and subscale scores. (see Supplementary Information for details). From the resultant data, 31 continuous measures indexing different psychological constructs were obtained from each participant. For all measures, baseline (8672 participants) and 2-year follow-up (6218 participants) data were included in the analyses, based on quality control of their MRI data and availability of their cognitive-psychological assessments (See Supplementary Information and Fig.S1 for more details).

### Clinical Diagnoses of Psychiatric Disorders

Participants were assessed at baseline and 2-year follow-up for the presence of current psychiatric diagnoses using the computerized version of the Kiddie Schedule for Affective Disorders and Schizophrenia for DSM-5 (KSADS-5) a psychometrically validated semi-structured psychiatric interview^80,90^. Psychiatric disorders (Depressive Disorder, Psychotic Disorder, Bipolar Disorder, Attention-deficit/hyperactivity Disorder, Substance Use Disorder, Alcohol Use Disorder, Obsessive-compulsive Disorder, Social Anxiety Disorder, Generalized Anxiety Disorder, Separation Anxiety Disorder, Eating Disorder, Conduct Disorder, Posttraumatic Stress Disorder, Oppositional Defiant Disorder, Specific Phobia, Panic Disorder, Agoraphobia, Disruptive Mood Dysregulation Disorder, Enuresis and Encopresis Disorder, and Tic Disorder) were used for the following analyses. For each specific disorder, the present diagnosis score was labeled “0” for absence of diagnosis and “1” for definitive diagnosis.

### Discovery of Brain–Cognition–Psychology Dimensions via Canonical Correlation Analysis (CCA)

To identify associations between the cortical structure of the youth brain (cortical area, volume, cortical thickness and sulcal depth) and cognitive-psychological assessments (Fig.1A) we conducted a CCA^91^ on the 2 datasets (see Supplementary Information and Fig. S2-S3 for more details). CCA identifies orthogonal latent variates (Fig.1B) from the brain and cognitive-psychological datasets while ensuring a maximized correlation between the 2 variates paired by their orders (modes). To identify meaningful brain-cognitive-psychological associations, we tested these modes for their statistical significance^92,93^, generalizability^41^, and redundancy index^38,41^ via permutation tests.

### Application of the Derived Pattern at Baseline to 2-Year Follow-up Measurements

To assess the identified association between the brain structural variate and individual differences in cognitive performance and psychological processes, we first derived a significant pattern from baseline brain and cognitive-psychological measurements. We then applied this pattern to the 2-year follow-up brain and cognitive-psychological measurements to compute the corresponding brain structural variate score and cognitive-psychological variate score at the later time point (see Supplement for more details). By subtracting the baseline variate scores from the 2-year follow-up scores, we obtained the brain structural variate score change and cognitive-psychological variate score changes over the 2-year period (see Fig. S3), thereby capturing how the same baseline-derived pattern evolved across time.

### Association Between the Brain Structural Variate, Clinical Diagnoses, and Status Transitions

To elucidate the clinical utility of the brain structural and cognitive-psychological variate in predicting psychiatric disorders, we investigated its associations with clinical diagnoses at both baseline and 2-year follow-up among participants categorized by KSADS diagnoses. Specifically, we examined the brain structural and behavioral measure under three conditions: (1) at baseline, (2) at the 2-year follow-up, and (3) the change between these two time points.

For disorder-specific, transdiagnostic spectrum and cumulative diagnostic-count comparisons, we first quantified the cross-sectional association between the brain variate and diagnostic status at each time point by relating baseline variate scores to baseline KSADS diagnoses and 2-year variate scores to follow-up KSADS diagnoses (see.1C). To probe the dynamics of these relationships, we then examined how changes in the variate score over two years covaried with diagnostic transitions (healthy-persistent group, disorder-new-onset group, disorder-remitted group and disorder-persistent group) (see Fig.1D) and additionally tested whether the baseline variate score predicted subsequent diagnostic shifts. This two-pronged approach allowed us to determine not only whether the variate tracks concurrent diagnostic status, but also whether individual differences in brain-behavior coupling confer risk or resilience to changes in psychiatric diagnosis over time. Further methodological details are provided in the Supplementary Information.

## Data availability

All data utilized in this study were from the ABCD Study (https://abcdstudy.org). Access to the data requires users to create an account through the National Institute of Mental Health (NIMH) Data Archive. After registration, users can follow the necessary procedures to obtain data access (https://data-archive.nimh.nih.gov/abcd).

## Acknowledgments

This research was supported by the Intramural Research Program of the National Institutes of Health (NIH). The contributions of the NIH author(s) were made as part of their official duties as NIH federal employees, are incompliance with agency policy requirements, and are considered Works of the United States Government. However, the findings and conclusions presented in this paper are those of the author(s) and do not necessarily reflect the views of the NIH or the U.S. Department of Health and Human Services.

